# Fast model-based ordination with copulas

**DOI:** 10.1101/2021.03.28.437086

**Authors:** Gordana C. Popovic, Francis K. C. Hui, David I. Warton

## Abstract

1. Visualising data is a vital part of analysis, allowing researchers to find patterns, and assess and communicate the results of statistical modeling. In ecology, visualisation is often challenging when there are many variables (often for different species or other taxonomic groups) and they are not normally distributed (often counts or presence-absence data). Ordination is a common and powerful way to overcome this hurdle by reducing data from many response variables to just two or three, to be easily plotted.
2. Ordination is traditionally done using dissimilarity-based methods, most commonly non-metric multidimensional scaling (nMDS). In the last decade however, model-based methods for unconstrained ordination have gained popularity. These are primarily based on latent variable models, with latent variables estimating the underlying, unobserved ecological gradients.
3. Despite some major benefits, a major drawback of model-based ordination methods is their speed, as they typically taking much longer to return a result than dissimilarity-based methods, especially for large sample sizes.
4. We introduce copula ordination, a new, scalable model-based approach to unconstrained ordination. This method has all the desirable properties of model-based ordination methods, with the added advantage that it is computationally far more efficient. In particular, simulations show copula ordination is an order of magnitude faster than current model-based methods, and can even be faster than nMDS for large sample sizes, while being able to produce similar ordination plots and trends as these methods.

## INTRODUCTION

Visualisation is a vital part of working with data in ecology, both for exploration, and to support conclusions from statistical analyses. Many studies in ecology however collect multivariate and discrete data, which are difficult to visualise. In this article, we focus on ordination (Zuur *et al*., 2007; Legendre & Legendre, 2012), a general and very commonly applied class of visualisation methods which aim to collapse or project multivariate data onto a small number (often two or three) dimensions in ways that preserve as much of the underlying structure as possible, such that it can be plotted to look for prevailing patterns. In this paper our focus is on ordination methods for multivariate abundance data e.g. species assemblages, although in principle the ideas discussed here would apply more generally to any multivariate analysis.

There are two broad categories of ordination methods in the current literature, traditional dissimilarity-based methods and model-based methods. Dissimilarity-based ordination methods begin by calculating a dissimilarity or distance matrix between sites (or more generally, observational units). Afterwards, these are collapsed to a small number of dimensions for plotting using an algorithm that attempts to preserve and display information about these relative distances, with the most widely used example of this being non-metric multidimensional scaling (MDS; Kruskal, 1964); see also van der Maaten & Hinton (2008) for a modern alternative. By contrast, model-based ordination methods explicitly model the underlying distribution of the response, using an extension of generalised linear mixed models known as latent variable models (or factor analytic models). As the name suggests, these models account for the correlation between taxa via the inclusion of a small number of latent variables, which are assumed to be random effects (Walker & Jackson, 2011; Hui *et al*., 2015; Warton *et al*., 2015). As they model the correlation between taxa, latent variables also act as natural ordination axes reflecting unobserved covariates. Both model-based and distance-based methods often give qualitatively similar ordinations e.g., see *Application to data* later on, as well as Hui *et al*. (2015), among others.

By directly modelling the data, model-based methods account for both the natural variation and the associated mean-variance relationships present in multivariate abundance data (Warton & Hui, 2017). Furthermore, we can use standard statistical tools to check the assumptions underlying the model, perform model selection, quantify the uncertainty in the estimated correlations between taxa, predict to existing and/or new sites, and incorporate extensions to (for instance) account for spatio-temporal correlations or imperfect detection (e.g., Warton *et al*., 2015; Ovaskainen *et al*., 2017; Hui *et al*., 2018; Tobler *et al*.,2019). All of this is generally more challenging to accomplish using dissimilarity-based ordination procedures.

One major drawback of most currently implemented model-based methods for ordination is that they tend to be considerably slower than dissimilarity-based methods, particularly when sample size (both the number of sites and/or taxa) is small to moderate. While there have been be some efforts to overcome this burden (e.g., Niku *et al*., 2019a; Tikhonov *et al*., 2020a), computation currently remains a major bottleneck when applying model-based approaches for ordination to large multivariate abundance datasets.

The primary reason why computation presents a major challenge for model-based approaches to ordination is that they tend to be built using hierarchical models with latent variables, formulated via a series of conditional distributions, from which it is difficult to calculate the marginal likelihoods or posterior distributions needed for estimation and inference (Walker & Jackson, 2011; Hui *et al*., 2015; Warton *et al*., 2015; Tikhonov *et al*., 2020b). To overcome this challenge, we present here an alternative family of multivariate models for ordination based on copulas. Copulas have only recently attracted interest in community ecology (Popovic *et al*., 2018; Anderson *et al*., 2019; Popovic *et al*., 2019). They can be considered as an extension of generalized linear models (GLMs, McCullagh & Nelder, 1989), and hence retain all the desirable statistical properties of model-based methods. Importantly, in contrast to hierarchical models, copulas are based on a *marginal* approach to modelling multivariate abundance data, so the relevant marginal likelihoods are much easier to compute, and the model is much faster to fit (see Fieberg *et al*., 2009; Muff *et al*., 2016, for examples of comparisons between conditional versus marginal approaches to modeling). In this paper, we propose a Gaussian copula model with a latent variable structure as a novel, fast model-based ordination method for community ecology. Simulations show our method is an order of magnitude faster than existing model-based methods, while producing largely similar ordination patterns and conclusions.

In the remainder of the article, we introduce copulas specifically for discrete data (given the prevalence of non-continuous responses in community ecology), and the proposed Gaussian copula latent variable model for ordination. The proposed method is implemented in the `R` package `ecoCopula`, via the `cord` function. We illustrate an example of copula-based ordination on an example multivariate abundance dataset, and conduct a simulation study to compare cord to existing model-based and distance based methods of ordination in terms of both accuracy and speed.

## MATERIALS AND METHODS

### Model-based methods for ordination

Multivariate abundance data, also known as multi-species or community composition data, consist of abundance measurements (generally presence/absence, counts, cover or biomass) simultaneously collected for a large number of taxa. Due to the discrete nature of such data, each taxon is typically modelled with a distribution appropriate to its characteristics *via* a generalised linear model (GLM; McCullagh & Nelder, 1989) or some variation thereof e.g., the negative binomial (Caraka *et al*., 2018) or zero-inflated count distribution for overdispered species counts, the binomial distribution (Golding *et al*., 2015) for presence-absences, and the Tweedie or Gamma distribution for biomass (Blakey *et al*., 2016) etc.

In order to model correlations between the taxa, as is central for ordination plots, it is necessary to construct a multivariate distribution for all taxa jointly. By far the most popular method of achieving this is via a hierarchical modelling framework, where latent variables (assumed to be Gaussian) form the ordination axes; see Appendix for mathematical details. However, estimation of a hierarchical model involves computationally burdensome steps, e.g., Markov Chain Monte Carlo sampling (in the case of Bayesian methods, Hui, 2021; Tikhonov *et al*., 2020b), or numerical integration of the marginal likelihood (in the case of likelihood-based methods, Niku *et al*., 2019b). In this paper, we consider an alternative model-based framework that is much faster to fit to multivariate abundance data – Gaussian copula models.

### Gaussian copula ordination

In this section, we introduce copulas as a means of constructing a joint distribution for multivariate abundance data, with a focus on ordination. Copulas are a flexible class of models which allow the modelling of data using any set of marginal distributions, coupled with the covariance structure of any desired multivariate distribution. The term “copula” comes from this idea of coupling a *marginal model* with a *multivariate model* that can handle covariance across response variables. To construct an ordination of multivariate abundances, we will couple marginal GLMs with a *factor analysis* for ordination, which we have implemented as the `cord` function in the `ecoCopula` package.

For the marginal model, we need to specify a separate model for each taxon that accounts for key properties of the data. A key feature of abundance data is its mean-variance relationship, we account for this using a GLM with an appropriate choice of distribution. This marginal model allows us to calculate cumulative probabilities for each taxon, we write as *F_j_*(·) the cumulative distribution function (or *CDF*) assumed for the *j*th taxon.

In a Gaussian copula model, abundances are mapped to copula values *z* that have a normal (or *Gaussian*) distribution, such that we can specify a multivariate model which assumes multivariate normality. For count data, and letting *y_ij_* denote the observed count for site *i* and taxon *j*, these copula values *z_ij_* satisfy

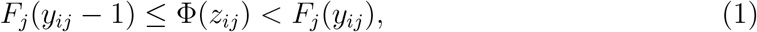

where *F_j_*(·) is the CDF assumed under the marginal GLM for the *j*th taxon, and Φ(·) is the CDF of the standard normal. More generally we can replace *F_j_*(*y_ij_*−1) in equation (1) with 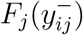, the left limit of *F_j_* at *y_ij_*. We note that these values are related to Dunn-Smyth residuals (Dunn & Smyth, 1996) which are often used for residual analysis in community ecology. Specifically, these residuals *r_ij_* satisfy the equation Φ(*r_ij_*) = *F*(*y*^−^)+*uf*(*y*), where *u* is a simulated value from the uniform distribution between zero and one, and *f*(·) is the probability density/mass function corresponding to the CDF *F*(·).

For the multivariate model, and hence ordination, we assume a factor analytic formulation, meaning we assume copula values *z_ij_*’s are multivariate normally distributed and satisfy

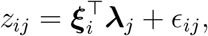

where ***ξ***_i_ denote the latent scores for site *i* and are assumed to be independent normal, **λ**_*j*_ are the corresponding factor loadings for taxon *j*, and *ϵ_ij_* are independent Gaussian errors, with variance 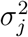 for taxon *j*. We construct an ordination of observations (sites) by plotting the estimated factor scores, and can construct a biplot by overlaying the estimated factor loadings, as is typically done in unconstrained ordination.

For unconstrained ordination, *F_j_*(·) is a GLM with only an intercept. To construct a residual ordination (Warton *et al*., 2015), *i.e*. ordination controlling for some measured variables, we can include these as predictors in the GLM.

We estimate the Gaussian copula ordination model by maximum likelihood, which in this case has no closed form due to the inequalities in equation (1). The maximisation employs an efficient iterative procedure where a factor analysis (we used the `factanal` function in `R`) is applied to iteratively reweighted Dunn-Smyth residuals. This makes the resulting procedure computationally efficient because the factor analysis likelihood has a closed form, and so the process of estimating the multivariate model is fast; see Appendix for details.

We conclude this section by highlighting out the two main assumptions required in specifying the Gaussian copula model: (1) the CDF *F_j_* has been specified correctly for all taxa *j* = 1,…, *D*; (2) the unobserved vector characterising the copula, ***ξ***_i_, comes from a multivariate Gaussian distribution. We require assumption (1) in order to be able to map from any set of variables to a set of Gaussian variables. But note that just because a set of variables is Gaussian does not necessarily mean they are multivariate Gaussian – hence the need for assumption (2), which essentially requires the further constraints of linearity and equal variance when looking at each copula variable as a function of other responses. Importantly, both these assumptions can be checked, say, by plotting Dunn-Smyth residuals and applying residual diagnostic tools.

## APPLICATION TO DATA

The hunting spider data is a well-known ecological dataset popularised in ter Braak (1986). It consists of counts of hunting spiders caught in pitfall traps, with *D* = 12 species found at *n* = 28 sites (van der Aart & Smeenk-Enserink, 1974). The data also contain six environmental variables thought to be associated with spider abundance, namely: dry soil mass; percent cover of bare sand; percent cover of fallen leaves or twigs; percent cover of moss; percent cover of herb layer and reflection of the soil surface with a cloudless sky. The primary question of interest in the original collection of the data was to identify the main environmental drivers of abundance of the species studied.

### Copula ordination

It is straightforward to carry out model-based ordination on the spider data using the `ecoCopula` package. Figure 1 presents an example of the estimated ordinations (unconstrained and residual) for these data. There are two key coding steps when constructing an ordination plot using the `ecoCopula` package: 1) fitting separate marginal models to each spider speces (to obtain the CDF *F_j_*), for which we provide the function `stackedsdm`; 2) then applying the function cord, which takes the output from the marginal model and returns the ordination to be plotted, assuming a Gaussian copula model.

**Figure 1:**
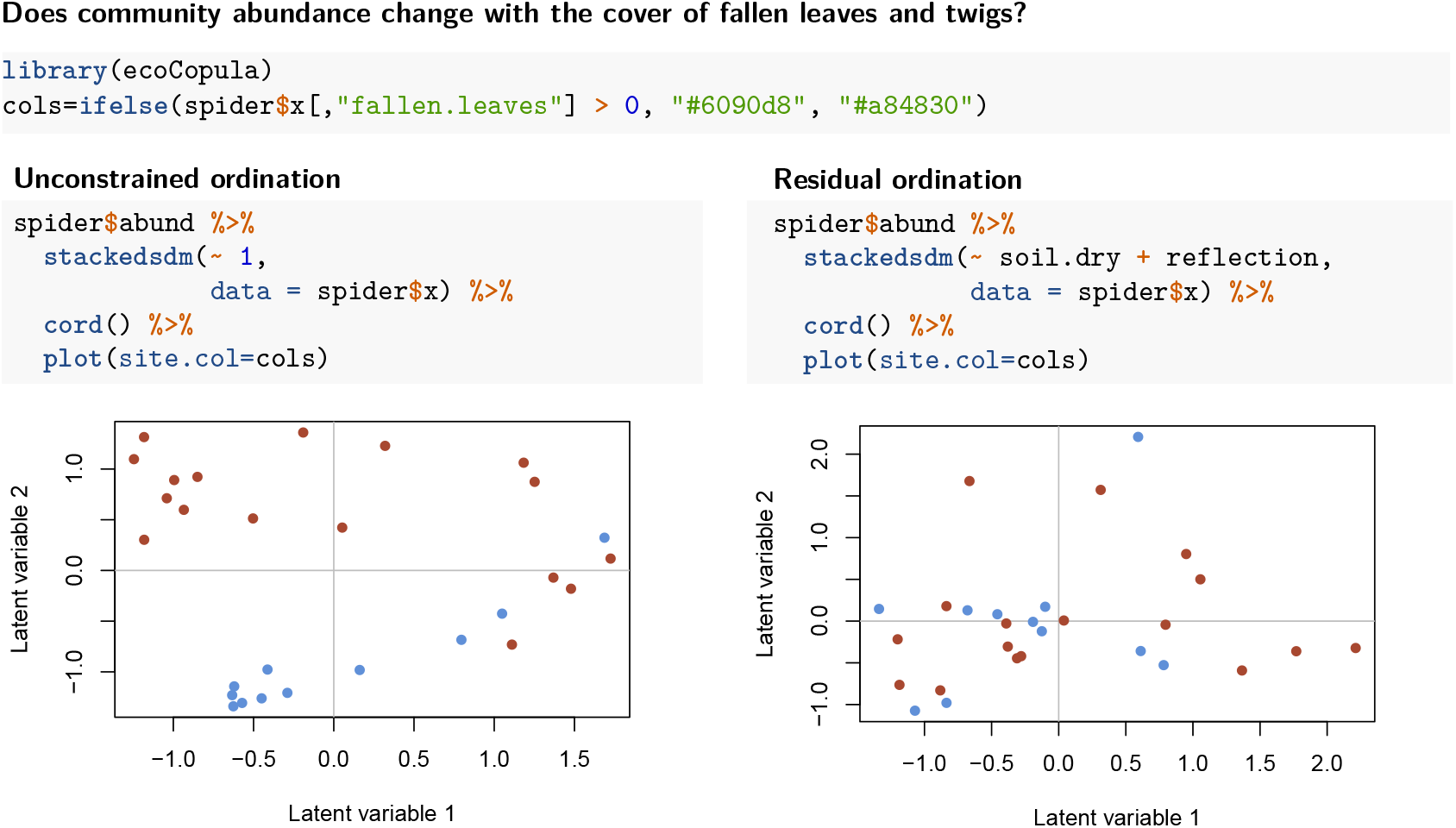
An illustration of estimated ordinations for the hunting spider data using the `ecoCopula` package. Sites are coloured according to presence (blue) or absence (red) of fallen leaves or twigs. In the unconstrained ordination (left), we see a clear separation of the sites based on this covariate, with sites with fallen leaves or twigs having predominantly negative values on the second ordination axis. On the right we have performed a residual ordination, controlling for moisture and reflectivity of the soil. The sites with and without fallen leaves and twigs are no longer well separated, indicating that moisture and reflectivity of the soil has explained some of the effect of fallen leaves and twigs on spider abundances.

In the first step, the `stackedsdm` function takes as input the matrix of multivariate abundances, a formula (which defaults to an intercept-only model, for an unconstrained ordination), and optionally a data frame storing covariates. Because it was written specifically for abundance data, it defaults to negative binomial regression if a family argument has not been entered, as does `manyglm` from the `mvabund` package (Wang *et al*., 2020), which could also be used with `cord`. The two main distinctions from `manyglm` are that `stackedsdm`: can take vector family input, which is useful if different response variables were collected in different ways; and it can take advantage of parallel processing to speed up fitting of the marginal models for very large datasets.

In the second step, the `cord` function applies the `factanal` function to iteratively reweighted Dunn-Smyth residuals from the marginal model fit. By default two factors are fitted, but other options can be specified via the `nlv` argument. The `cord` function was written so it could be applied to marginal model fits constructed using `stackedsdm` or `manyglm`. In principle, other functions could be used to fit the marginal models too, as long as they have a `residuals` function and store fitted values as a matrix response.

### Competing ordination methods

There are several competing methods commonly used for ordination in community ecology. Hierarchical model-based ordination methods e.g., `R` packages `Hmsc` (Tikhonov *et al*.,2020c), `boral`, and `gllvm` use latent variables to define the ordination axes. For ordination purposes, these all effectively fit the same type of model but by slightly different means of estimation. Not surprisingly then, speed is arguably the main distinguishing feature between these approaches, with the Bayesian methods being slower than their likelihood-based counterparts. From the literature and our experience, we found that `gllvm` was the fastest existing package that implemented a hierarchical approach to ordination. For example, carrying out a model-based unconstrained ordination on the spider data with `gllvm` takes 1.06 seconds, compared to 25.82 seconds for `boral` and 79.48 seconds for `Hmsc` (each with 10000 MCMC samples).

There are many ordination methods in common use in community ecology that do not use a model-based framework, with nMDS being the most popular at the time of writing. We include this in our comparison (using the `metaMDS` function in the `vegan` package, applied after fourth root transformation using the Bray-Curtis dissimilarity). In addition, we considered another popular approach in detrended correspondence analysis (DCA; Hill & Gauch, 1980), as implemented using the `decorana` function in the `vegan` package (Oksanen *et al*., 2019).

Applying the range of ordination methods discussed above to the spider data, we see that they give qualitatively similar results (e.g., in Fig 2), despite using quite different formulations.

**Figure 2:**
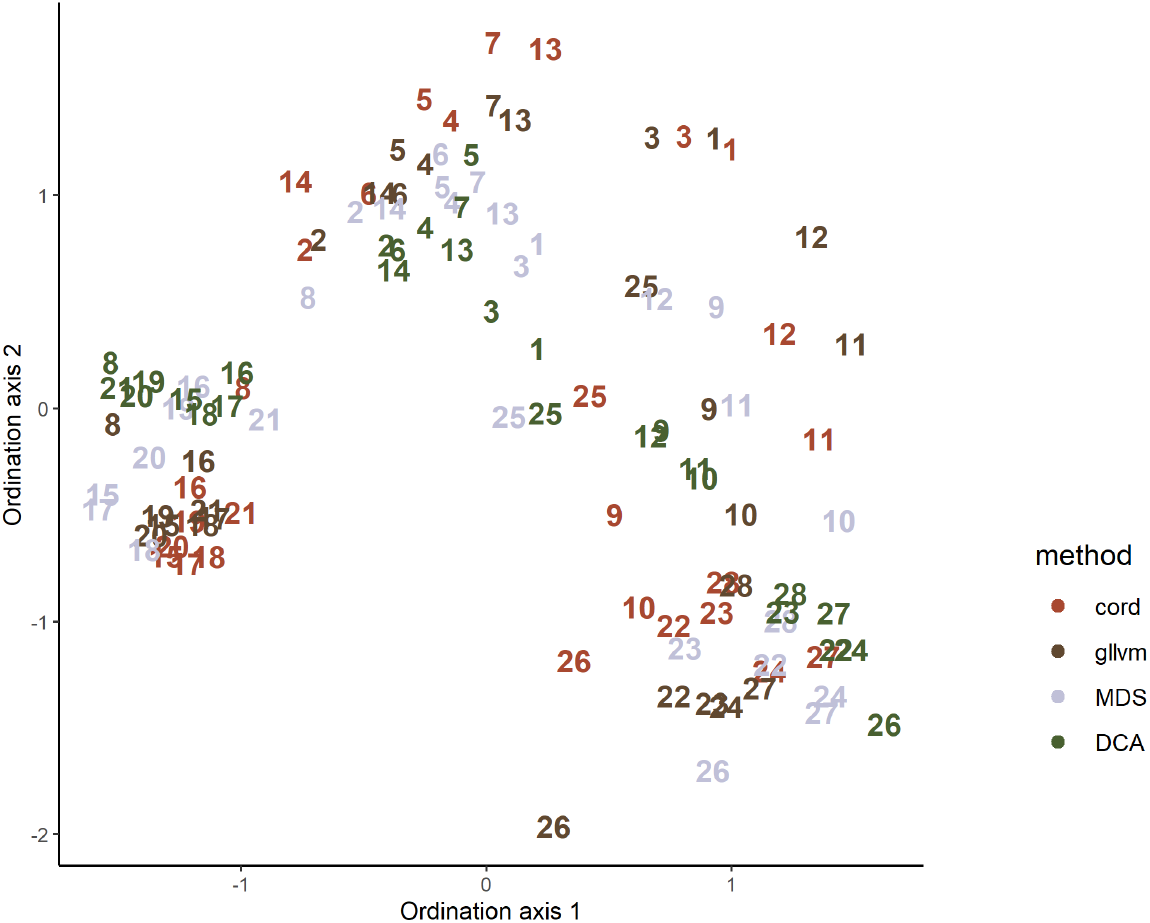
Unconstrained ordination scores estimated on the cord function in the `ecoCopula` package (which uses the proposed Gaussian copula method, red), the `gllvm` package (which is computationally the most efficient package for implementing ordination via hierarchical modeling currently available, brown), nMDS (based on applying the fourth root transformation and the Bray-Curtis dissimilarity, grey), and DCA (green). All scores have been rotated and scaled using `vegan::procrustes`. Qualitatively, it can be seen that these ordinations are fairly similar, despite being estimate using very different approaches.

**Figure 3:**
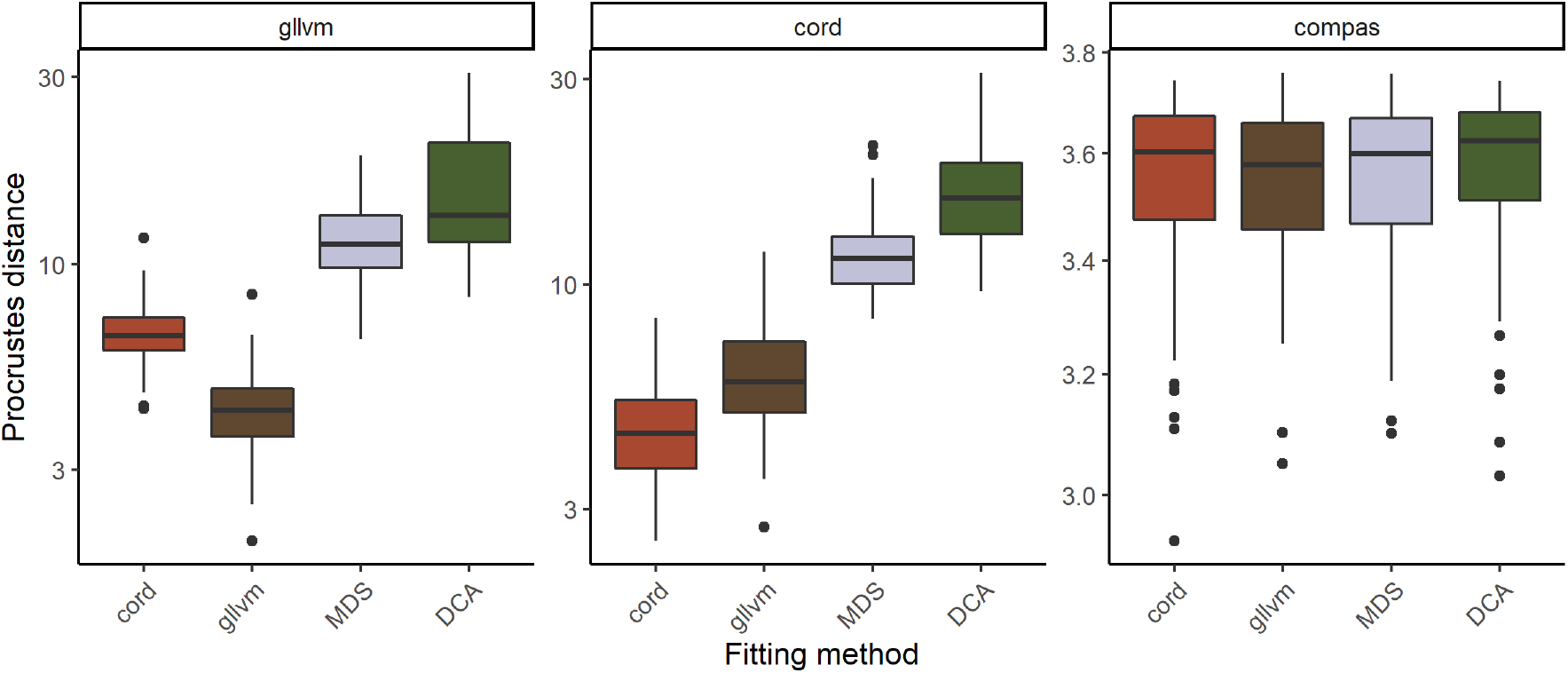
Comparative boxplots assessing ordination score recovery of the four ordination methods fitted to data generated from a hierarchical modelling approach (`gllvm`), the proposed Gaussian copula model (`cord`), and a more “neutral” approach (`compas`). Results showed that `gllvm` and `cord` performed best when data were generated from their respective models, while for `compas` all four methods performed similarly.

## SIMULATION STUDY

To quantitatively compare the Gaussian copula ordination method to existing methods in terms of speed and accuracy, we conducted a simulation study using the spider data as the basis for simulating multivariate abundance data.

### Simulation design

For assessing ordination score recovery, we simulated datasets that mimicked properties of the hunting spider data, using one of three approaches: 1) the `simulate` function in the `gllvm` package, which simulates abundances from a latent variable model i.e, a hierarchical modelling approach; 2) the `simulate` function for `cord` models in the `ecoCopula` package, which is based on our proposed Gaussian copula model, and; 3) `compas` (Minchin, 1987), which simulates multivariate count data based on generating unimodal curves representing species responses to one or more indirect gradients. To simulate from `gllvm` and `cord`, we fit the corresponding model to the spider data without covariates, and then simulate from the resulting model fit treating the estimated parameters and predicted latent variables as the true values. To simulate using `compas`, we use the scores obtained from applying nMDS to the spider data (assuming a fourth root transformation and Bray-Curtis dissimilarity), and then apply the `compas` function from the `CommEcol` package (Melo, 2019), and using default values for all arguments as appropriate. For each case, we simulated 100 multivariate count datasets.

For each simulated dataset, we then compared the performance of the estimated ordination scores using four competing methods: `gllvm`, `cord`, nMDS, and DCA. Performance was assessed using the Procrustes distance between the true and estimated ordination scores. Note we did not consider other packages such as `Hmsc` or `boral`, as we found they tended to produce similar answers to `gllvm`, but, as is the case above in *Competing ordination methods* took considerably longer to produce the ordination.

Next, as an assessment of computation time between the methods, we simulated new multivariate count datasets with varying numbers of sites and/or taxa. We then fitted the same four ordination methods as above to obtain ordination scores, tracking computation time in seconds required for each method. For each combination of number of sites *n* and number of taxa *D*, we simulated 20 datasets, and averaged computation time across the 20 fits for each method. Note that because computation speed rather than ordination accuracy was of interest at this point, multivariate abundance data were simulated from the `gllvm` simulation model only.

### Results

#### Ordination score recovery

As expected, the precise performance of each method, while qualitatively similar (Fig 2), depended strongly on how the data were generated. When data were simulated using `gllvm` i.e., a latent variable model, then `gllvm` performed best at recovering ordination scores, followed by the proposed Gaussian copula method using `cord`. Similarly, when the data were simulated from a Gaussian copula model, `cord` performed best at recovering the scores followed by the hierarchical modelling approach using `gllvm`. When data were simulated using `compas`, none of the methods stood out in terms of ordination recovery. The two model-based approaches to ordination clearly outperformed nMDS and DCA when the data generation process was model-based (even if it was the incorrect model), but when the data generation process was more “neutral” i.e., `compas`, there was no clear difference between the four approaches.

#### Computation speed

In all simulations, the proposed Gaussian copula method i.e., `cord`, was noticeably faster than `gllvm` (Fig 4), and indeed we anticipate that as an alternative approach to ordination it will likely be faster than any other currently available software that uses a hierarchical/latent variable approach for ordination. For a small number of sites *n*, `cord` was slower than the dissimilarity-based methods we considered. Interestingly though, `cord` scaled better with the number of sites *n* compared to other methods, and for large enough *n* e.g., approximately *n* ≥ 200 it was faster than nMDS (under its default settings on `vegan`, Fig 4). Dissimilarity-based methods were consistently the fastest methods when the number of taxa *D* became large.

**Figure 4:**
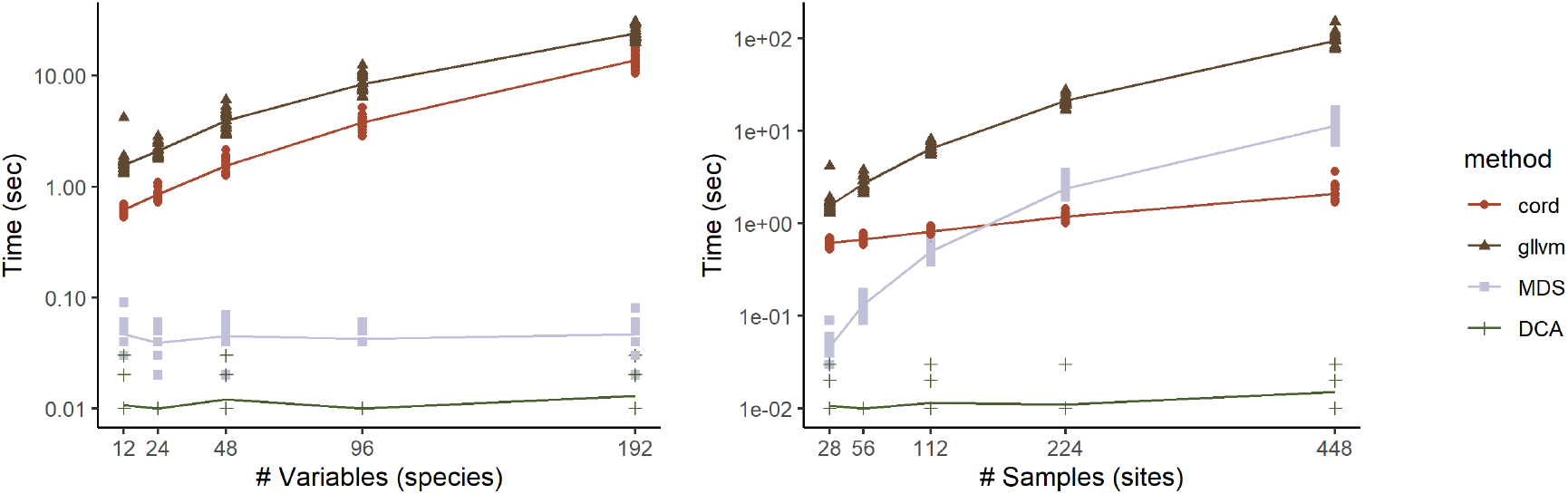
Comparison of computation times when constructing an ordination using different approaches. The proposed Gaussian copula methods `cord` was noticeably faster than the fastest model-based competitor `gllvm`, and also faster than nMDS for large values of *n*. Dissimilarity-based methods were faster when the number of taxa *D* was large and/or the number of sites *n* was small.

## DISCUSSION

In this article, we have proposed a computationally efficient approach to model-based ordination which performs well at recovering ordination scores in community ecology, and is faster and more scalable than all existing model-based ordination methods in the literature. Interestingly, it was even faster than the popular dissimilarity-based method nMDS for datasets with a moderate number of taxa and a moderate to large number of sites. Two other key features of the proposed approach include the straightforward capability to consider a large range of data distributions for the responses, to have different distributions for different taxa i.e., multivariate abundance data with mixed responses, and the possibility of residual ordination, *i.e*. controlling for environmental factors in the ordination. The proposed Gaussian copula method is implemented in the `R` package `ecoCopula`, which is now available on `CRAN`. The package also contains a `simulate` function, which allows users to simulate data with similar characteristics to the observed data assuming a Gaussian copula model; this is useful for sample size calculations for study design, diagnostics, and visual inference, among other potential applications.

The speed of the copula approach to ordination, relative to hierarchical models, is driven by the use of marginal models, and the choice of a Gaussian distribution for the multivariate model. This meant that the latent variable model was estimated *via* a Gaussian likelihood function, which is generally well-behaved and quick to compute (largely because for Gaussian likelihoods, we can work directly with the sample covariance matrix rather than the full dataset). While we only explored this here in the context of ordination, we expect this to be generally true across other covariance models (such as graphical modelling; Popovic *et al*., 2019), and anticipate a wealth of potential extensions and applications of Gaussian copula models in ecology.

For example, further ordination functionality could include an extension of redundancy analysis to handle data from any marginal distribution, since this can be understood as a form of reduced rank regression on multivariate Gaussian data (Bach & Jordan, 2005; Stoica & Viberg, 1996), and therefore can be applied to the estimated copula model using a similar estimation algorithm to that used in `ecoCopula` (Popovic *et al*., 2018). Fast model-based ordination can also be combined with high-dimensional data visualisation and visual inference tools such as those implemented in the `tourr` package (Wickham *et al*., 2011) to visualise three or more latent variables, and we are currently undertaking research on this front. In addition, marginal models can be extended in a variety of ways, from modelling non-linear relationships using generalised additive models (Wood, 2017), to accounting for detection (Tobler *et al*., 2019), other response types (e.g., plant cover data, Damgaard *et al*., 2020), and explaining changes in abundance with traits via a fourth-corner model (Brown *et al*., 2014). Spatially and/or temporally indexed multivariate abundance data can be modelled using more structured latent scores in the Gaussian copula model (*e.g*. Thorson *et al*., 2015, 2016); this can be interpreted as a set of unobserved and spatially smoothed environmental predictions or ordination axes.

It should be noted that, along of the lines of what has been discussed in the article, handling such spatial and/or temporal multivariate abundance data though a copula model is expected to be computationally far more efficient and scalable than hierarchical or latent variable approaches (Hui *et al*., In press), since it can be straightforwardly done through the covariance of *Z_ij_*’s while maintaining the multivariate normality assumption. Finally, copula models can be used as a basis for performing likelihood-based inference about multivariate abundance data, for example to test for effects of a one or more treatment or environmental covariates. In summary, the copula-based framework has considerable potential in community ecology, due to both its flexibility and its advantages in computational efficiency and scalability, and we look forward to seeing how this literature develops in the future.

## ACKNOWLEDGEMENTS

GCP was supported by the Australia Postgraduate Award and ARC Discovery Project project (DP180103543) awarded to DIW. FKCH was supported an ARC Discovery Early Career Research Fellowship (DE200100435).

## AUTHOR CONTRIBUTIONS

GCP led the writing and analysis. All authors conceived and developed ideas, contributed critically to the manuscript editing, and gave final approval for publication.

## DATA ACCESSIBILITY

The hunting spider data are available as part of the `ecoCopula` package.

## APPENDIX

### Ordination with hierarchical models

Let ***y*** be the response matrix with elements *y_ij_* for site *i* = 1, 2,…, *N* and taxon *j* = 1, 2,…, *D*. Then a typical hierarchical modelling approach to ordination models the conditional mean of each element, denoted as *μ_ij_*, via an extension of GLMs, where a small number *d* ≪ *D* of latent variables ***ξ**_i_* and an (optional) error *ϵ_ij_* are included on the linear predictor scale,

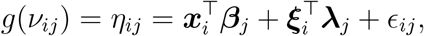

where ***x_i_*** is a vector of observed predictors for site *i*, ***β**_j_* are the corresponding taxon-specific coefficients, ***ξ**_i_* denote the latent variables, **λ**_*j*_ are the corresponding factor loadings. The quantity *g*(·) denotes some known link function relating the mean to the linear predictor. If included, the error terms are assumed to be independent across responses, such that the variance-covariance matrix **Ψ** of the vector ***ϵ**_i_* = (*ϵ*_*i*1_,…, *ϵ*_*iD*_) is diagonal. On the linear predictor scale, the covariance and hence correlation between the taxa is driven by the factor loadings. In particular, if we define ***η**_i_* = (*η*_*i*1_,…, *η_iD_*), then the variance-covariance at site *i* is then given by Cov(***η**_i_*) = **Σ** = **ΛΛ**^T^ + **Ψ**, where **Λ** is the loadings matrix formed by stacking the **λ**_*j*_’s as row vectors. The hierarchical nature of the latent variable model derives from the fact that it can be formulated conditionally. That is, we first assume a distribution for the latent variables e.g., the elements of **xi_*i*_** are assumed to follow a standard Gaussian distribution. Then, conditionally on the latent variables, we assume a distribution for the responses as given by the GLM framework, where the mean of the response is related to the measured covariates, latent variables, and errors (if appropriate) as defined above.

With regards to ordination, from the above specification we see that the latent variables can be understood as modelling unmeasured environmental gradients, while the factor loadings quantify the response of each taxon to each of these underlying gradients (similar to regression coefficients for a measured covariate). If the number of latent variables *d* is small (typically *d* = 2 or 3), then the estimated loadings and latent scores can be plotted separately, or can be combined into a *biplot* to visualise relationships between different sites, and between different taxa and between taxa and sites.

Depending on what additional information is available, there are a wealth of extensions that could be made to the hierarchical latent variable framework to further the ordination analysis e.g., the latent variables could be spatially and/or temporally varying (Thorson *et al*., 2015), and/or included at different scales (Björk *et al*., 2018). However for simplicity, we will focus on the specification as given above, noting that the proposed approach to model-based ordination via copulas detailed below may also be extended in such directions as avenues of future research.

### Copulas for multivariate abundance data

#### Copulas

Copula modelling is rooted in Sklar’s Theorem (Sklar, 1959), which shows that any multivariate distribution can be written as a copula model, as long as the marginal and copula distributions are appropriately chosen. From a modelling perspective, the main appeal of copulas is that the covariance structure can be specified independently of the marginal distributions, making them very flexible. In addition, their marginal specification makes them computationally efficient, fast to fit, and readily parallelizable.

In more detail, copulas can be viewed as providing a means of starting from a common non-Gaussian data distributions (Poisson, negative binomial, Tweedie, *etc*), and then extending them to multiple, *correlated* responses in a flexible manner. The idea is to map from any data distribution to a convenient copula family that facilitates multivariate modelling, most commonly, the multivariate Gaussian distribution as is done in our main text with Gaussian copula method. This mapping occurs using the probability integral transform (PIT), a basic property of continuous probability distributions which allows observed data values *y* to be converted to uniformly distributed values between zero and one via its corresponding cumulative distribution function (CDF), *F*(*y*). The PIT hence provides a way to transform between any two continuous distributions, in our case, between the desired data distribution and the copula distribution, by going though the standard uniform distribution itself. The latter copula distribution is where the multivariate properties of the data, including covariances and correlations, are straightforwardly formulated and estimated. We highlight that while there are a wide variety of possible choices of copula distributions (see Nelsen, 2007, for an introduction to copulas), in our case we will focus on the using the multivariate Gaussian distribution as the copula distribution, given its simplicity and relative easy of interpretation in the context of community ecology.

#### Discrete Gaussian copulas

To construct a copula model, we require a set of marginal distributions, one for each taxon, along with a multivariate (copula) distribution to model the correlations between taxa. As ecologists often have discrete data, the marginal distributions *F_j_*(· are, like in the hierarchical case, chosen to account for key properties of the data e.g., using a negative binomial or zero-inflated count GLM for overdispersed counts. Note however that when the marginal distributions are discrete, the PIT properties do not strictly hold, in the sense that the transformation is no longer a unique mapping. Nevertheless, we can still use this concept to derive a discrete Gaussian copula with marginal distributions *F_j_*; *j* = 1,…, *D* as a function an unobserved multivariate Gaussian variable.

To understand how this discrete Gaussian copula can be formulated, it is useful to first recap the idea of a randomized quantile or Dunn-Smyth residual (Dunn & Smyth, 1996) as discussed in the main text. The Dunn-Smyth residual is illustrated in the left panel of Fig 5. Specifically, instead of calculating *F*(*y*) as in the PIT, we first simulate a random value between *F*(*y*^−^) and *F*(*y*) where *y*^−^ generically denotes the previous observable value of a response *y* e.g., for counts these must occur at integer values, and then transform this value using the inverse CDF of a standard normal random variable. Mathematically, this can be written as 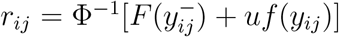, where *u* is a simulated value from the uniform distribution between zero and one and *f*(·) is the corresponding density function. For discrete data *y*, it can be subsequently shown that the Dunn-Smyth residual *r_ij_* follows a standard univariate Gaussian distribution, provided the CDF *F*(*y*) has been specified correctly. Dunn-Smyth residuals have been growing in popularity in ecology (Warton *et al*., 2017) and are used in popular packages such as `mvabund` (Wang *et al*., 2020), `boral` (Hui, 2021), `gllvm` (Niku *et al*., 2019b) and `DHARMa` (Hartig, 2020), largely for the purposes of residual analysis. Here however, we use the the idea of a Dunn-Smyth residual to map from a discrete distribution (Fig 5, left panel), through the uniform distribution (centre panel), to the copula distribution of our choice (right panel). We note here that for each value of *y*, there are many possible values of *r*, however for each *r*, there is only one value of the response *y* that could have produced it. This is a consequence of the fact that, as remarked previously, the PIT is no longer a unique mapping for discrete data.

**Figure 5:**
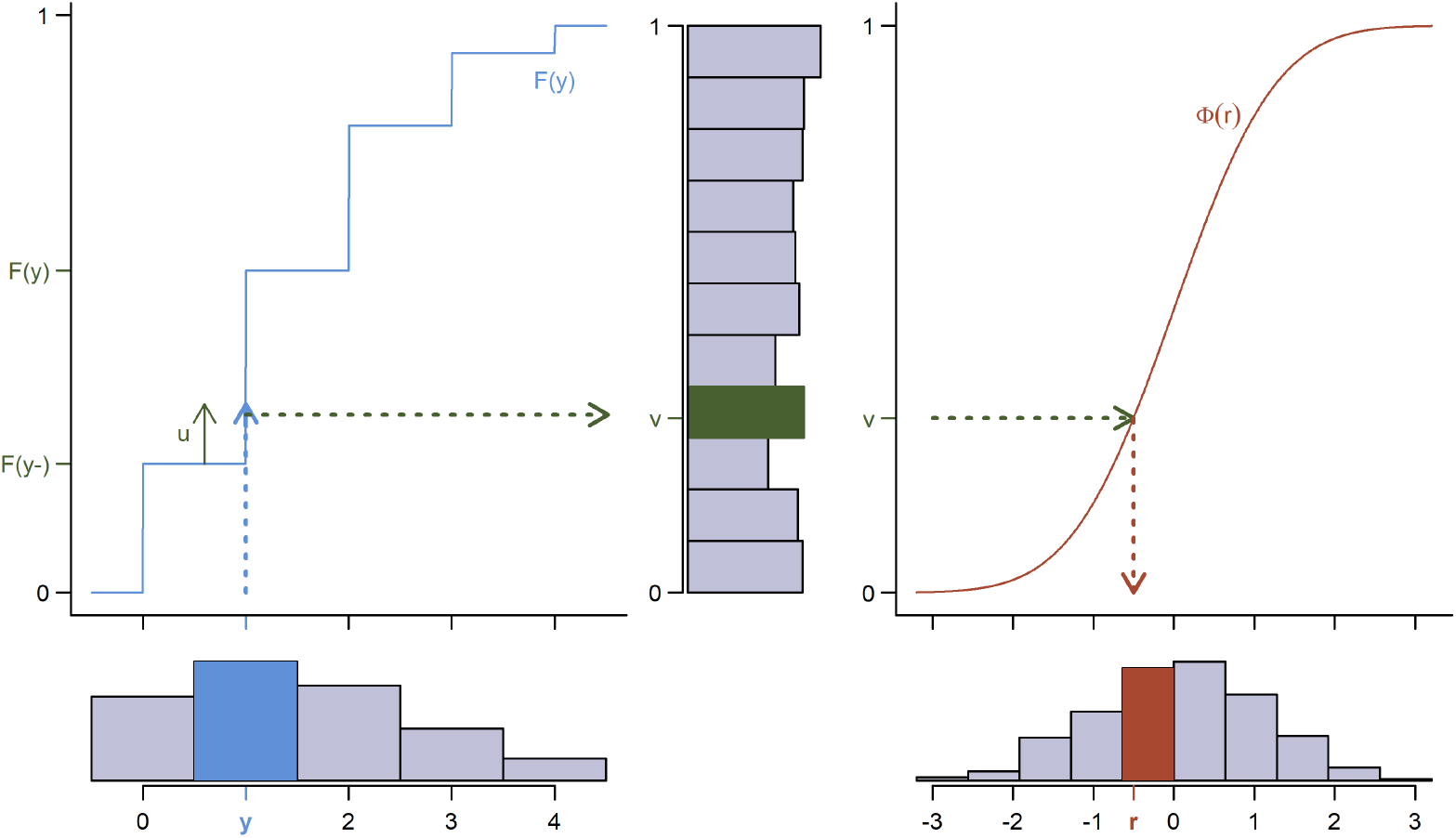
To generate Dunn-Smyth residuals *r* for a discrete observation *y* with CDF *F*(*y*) (left panel), we first simulate a uniform value u, (green) between *F*(*y*^−^) and *F*(*y*). The resulting *v* = *F*(*y*^−^) + *u* is uniform between zero and one (centre panel). We then transform v though the inverse of the Gaussian CDF, *r* =Φ^−1^(*v*).

Based on the idea of Dunn-Smyth residuals, we now discuss how to fit a Gaussian copula model to discrete data. For site *i* = 1,…, *n*, let ***z**_i_* = (*z*_*i*1_, *z*_*i*2_,…, *z*_*iD*_) denote a *D*-dimensional unobserved multivariate vector, arising from the copula distribution *f*(*z_i_*). Then the conditional distribution of *y_ij_* given ***z**_i_* is equal to one when 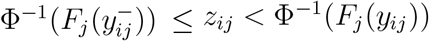, and zero otherwise. In other words, if we know *z_ij_* then we also know *y_ij_*, as it must be equal to whichever the value of the response that could have produced *z_ij_*, analogous to the idea of Dunn-Smyth residuals above. More formally,

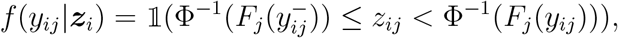

where 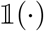 denotes the indicator function. To find the marginal distribution of **y**_*i*_, we first write the joint distribution of **y**_*i*_ and ***z***,

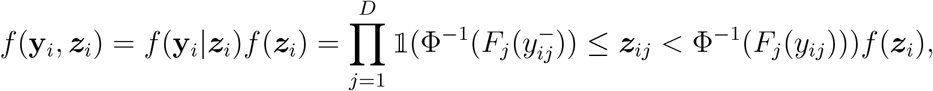

and then integrate out the unobserved vector ***z**_i_*, as follows

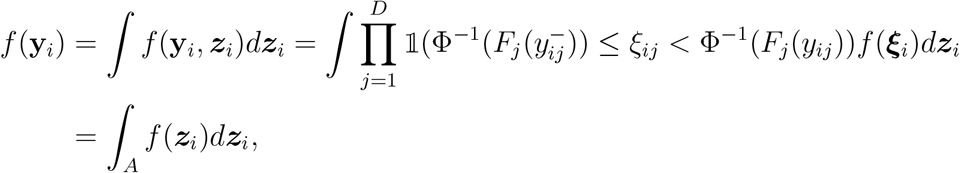

where 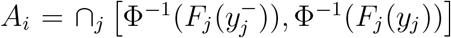. That is, the marginal likelihood of the data at site *i* is constructed by integrating the copula distribution *f*(***z***) over the region *A*, a hypercube with boundaries defined by the PIT.

By choosing *f*(***z**_i_*) to follow a multivariate Gaussian distribution, the above marginal likelihood then formally defines the Gaussian copula model. In practice, to fit such a model we want to first develop an approach by which to estimate this likelihood function (given data collected at a set of *n* sites), and then proceed find the values of model parameters (e.g., coefficients ***β**_j_*, loadings **λ**_*j*_, **Ψ** and so on) that maximise *f*(***y**_i_*). The main challenge lies in the former i.e., to approximate the integral. Perhaps not surprisingly, it turns out that we can use the idea of Dunn-Smyth residuals to estimate *f*(***y**_i_*) via importance sampling Popovic *et al*. (see 2018, for details). Essentially, we construct many sets of Dunn-Smyth residuals, and then take a weighted average of the likelihood evaluated at these residual values, weighted by how well each residual fits the assumed multivariate Gaussian copula distribution. Critically, the resulting approximate likelihood then has the form of a weighted sum of Gaussian likelihood terms, so maximum likelihood estimation of (all the parameters in) the copula model becomes relatively straightforward by adopting standard algorithms for estimating Gaussian distributed data; we refer the reader to Popovic *et al*. (2018) for the mathematical details underlying this, noting that we have implemented this in the `R` package `ecoCopula`. As mentioned previously, one major advantage of this estimation procedure, and of the copula model as a whole, is its computationally efficiency and scalability relative to estimating hierarchical models. This largely results from the fact that, by effectively transforming the problem from working with data from an complex high-dimensional multivariate discrete distribution to working with Dunn-Smyth residuals from a multivariate Gaussian distribution, we can use the many well-known properties and tools of the latter to perform likelihood-based estimation and inference. For example, we can use the sample covariance matrix (noting sampling from a multivariate Gaussian distribution is relatively straightforward) as a sufficient statistic for the correlation matrix of the copula distribution. In turn, we avoid the high-dimensional integrals inherent in estimating hierarchical or conditional models, instead replacing it with the need to sample from a tractable copula distribution, which in the case of a Gaussian copula model can be very efficiently.

## Notes

### Competing Interest Statement

The authors have declared no competing interest.

## References

Anderson, M. J., de Valpine, P., Punnett, A. & Miller, A. E. (2019). A pathway for multivariate analysis of ecological communities using copulas. Ecology and Evolution, 9, 3276–3294.

Bach, F. R. & Jordan, M. I. (2005). A Probabilistic Interpretation of Canonical Correlation Analysis. Tech. Rep. 688, Department of Statistics, University of California, Berkeley.

Björk, J. R., Hui, F. K. C., O’Hara, R. B. & Montoya, J. M. (2018). Uncovering the drivers of host-associated microbiota with joint species distribution modelling. Molecular ecology, 27, 2714–2724.

Blakey, R. V., Law, B. S., Kingsford, R. T., Stoklosa, J., Tap, P. & Williamson, K. (2016). Bat communities respond positively to large-scale thinning of forest regrowth. Journal of Applied Ecology, 53, 1694–1703.

Brown, A. M., Warton, D. I., Andrew, N. R., Binns, M., Cassis, G. & Gibb, H. (2014). The fourth-corner solution–using predictive models to understand how species traits interact with the environment. Methods in Ecology and Evolution, 5, 344–352.

Caraka, R. E., Shohaimi, S., Kurniawan, I. D., Herliansyah, R., Budiarto, A., Sari, S. P. & Pardamean, B. (2018). Ecological show cave and wild cave: negative binomial gllvm’s arthropod community modelling. Procedia Computer Science, 135, 377–384.

Damgaard, C., Hansen, R. R. & Hui, F. K. (2020). Model-based ordination of pin-point cover data: Effect of management on dry heathland. Ecological Informatics, 60, 101155.

Dunn, P. K. & Smyth, G. K. (1996). Randomized quantile residuals. Journal of Computational and Graphical Statistics, 5, 236–244.

Fieberg, J., Rieger, R. H., Zicus, M. C. & Schildcrout, J. S. (2009). Regression modelling of correlated data in ecology: subject-specific and population averaged response patterns. Journal of Applied Ecology, 46, 1018–1025.

Golding, N., Nunn, M. A. & Purse, B. V. (2015). Identifying biotic interactions which drive the spatial distribution of a mosquito community. Parasites & vectors, 8, 367.

Hartig, F. (2020). DHARMa: Residual Diagnostics for Hierarchical (Multi-Level / Mixed) Regression Models.

Hill, M. O. & Gauch, H. G. (1980). Detrended correspondence analysis: an improved ordination technique. Classification and ordination, pp. 47–58. Springer.

Hui, F. K. C. (2021). boral: Bayesian Ordination and Regression AnaLysis. R package version 2.0.

Hui, F. K. C., Hill, N. A. & Welsh, A. H. (In press). Assuming independence in spatial latent variable models: Consequences and implications of misspecification. Biometrics.

Hui, F. K. C., Sara, T., Pledger, S., Foster, S. D. & Warton, D. I. (2015). Model-based approaches to unconstrained ordination. Methods in Ecology and Evolution, 6, 399–411.

Hui, F. K. C., Tanaka, E. & Warton, D. I. (2018). Order selection and sparsity in latent variable models via the ordered factor LASSO. Biometrics, 74, 1311–1319.

Kruskal, J. B. (1964). Multidimensional scaling by optimizing goodness of fit to a non-metric hypothesis. Psychometrika, 29, 1–27.

Legendre, P. & Legendre, L. (2012). Numerical ecology. Elsevier.

McCullagh, P. & Nelder, J. A. (1989). Generalized Linear Models, Volume 37. CRC Press.

Melo, A. S. (2019). CommEcol: Community Ecology Analyses.

Minchin, P. R. (1987). Simulation of multidimensional community patterns: towards a comprehensive model. Vegetatio, 71, 145–156.

Muff, S., Held, L. & Keller, L. F. (2016). Marginal or conditional regression models for correlated non-normal data? Methods in Ecology and Evolution, 7, 1514–1524.

Nelsen, R. B. (2007). An introduction to copulas. Springer Science & Business Media.

Niku, J., Brooks, W., Herliansyah, R., Hui, F. K. C., Sara, T. & Warton, D. I. (2019a). Efficient estimation of generalized linear latent variable models. PloS one, 14, e0216129.

Niku, J., Hui, F. K. C., Taskinen, S. & Warton, D. I. (2019b). gllvm: Fast analysis of multivariate abundance data with generalized linear latent variable models in R. Methods in Ecology and Evolution, 10, 2173–2182.

Oksanen, J., Blanchet, F. G., Friendly, M., Kindt, R., Legendre, P., McGlinn, D., Minchin, P. R., O’Hara, R. B., Simpson, G. L., Solymos, P., Stevens, M. H. H., Szoecs, E. & Wagner, H. (2019). vegan: Community Ecology Package.

Ovaskainen, O., Tikhonov, G., Norberg, A., Guillaume Blanchet, F., Duan, L., Dunson, D., Roslin, T. & Abrego, N. (2017). How to make more out of community data? A conceptual framework and its implementation as models and software. Ecology letters, 20, 561–576.

Popovic, G. C., Hui, F. K. C. & Warton, D. I. (2018). A general algorithm for covariance modeling of discrete data. Journal of Multivariate Analysis, 165, 86 – 100.

Popovic, G. C., Warton, D. I., Thomson, F. J., Hui, F. K. C. & Moles, A. T. (2019). Untangling direct species associations from indirect mediator species effects with graphical models. Methods in Ecology and Evolution, 10, 1571–1583.

Sklar, M. (1959). Fonctions de répartition à n dimensions et leurs marges. Université Paris.

Stoica, P. & Viberg, M. (1996). Maximum likelihood parameter and rank estimation in reduced-rank multivariate linear regressions. IEEE Transactions on Signal Processing, 44, 3069–3078.

ter Braak, C. J. F. (1986). Canonical Correspondence Analysis: A New Eigenvector Technique for Multivariate Direct Gradient Analysis. Ecology, 67, 1167–1179.

Thorson, J. T., Ianelli, J. N., Larsen, E. A., Ries, L., Scheuerell, M. D., Szuwalski, C. & Zipkin, E. F. (2016). Joint dynamic species distribution models: a tool for community ordination and spatio-temporal monitoring. Global Ecology and Biogeography, 25, 1144–1158.

Thorson, J. T., Scheuerell, M. D., Shelton, A. O., See, K. E., Skaug, H. J. & Kristensen, K. (2015). Spatial factor analysis: a new tool for estimating joint species distributions and correlations in species range. Methods in Ecology and Evolution, 6, 627–637.

Tikhonov, G., Duan, L., Abrego, N., Newell, G., White, M., Dunson, D. & Ovaskainen, O. (2020a). Computationally efficient joint species distribution modeling of big spatial data. Ecology, 101, e02929.

Tikhonov, G., Opedal, Ø. H., Abrego, N., Lehikoinen, A., de Jonge, M. M. J., Oksanen, J. & Ovaskainen, O. (2020b). Joint species distribution modelling with the R-package Hmsc. Methods in Ecology and Evolution, 11, 442–447.

Tikhonov, G., Ovaskainen, O., Oksanen, J., de Jonge, M., Opedal, O. & Dallas, T. (2020c). Hmsc: Hierarchical Model of Species Communities.

Tobler, M. W., Kéry, M., Hui, F. K. C., Guillera-Arroita, G., Knaus, P. & Sattler, T. (2019). Joint species distribution models with species correlations and imperfect detection. Ecology, 100, e02754.

van der Aart, P. & Smeenk-Enserink, N. (1974). Correlations Between Distributions of Hunting Spiders (Lycosidae, Ctenidae) and Environmental Characteristics in a Dune Area. Netherlands Journal of Zoology, 25, 1–45.

van der Maaten, L. & Hinton, G. (2008). Visualizing Data using t-SNE. Journal of Machine Learning Research, 9, 2579–2605.

Walker, S. C. & Jackson, D. A. (2011). Random-effects ordination: describing and predicting multivariate correlations and co-occurrences. Ecological Monographs, 81, 635–663.

Wang, Y., Naumann, U., Eddelbuettel, D., Wilshire, J. & Warton, D. (2020). mvabund: Statistical Methods for Analysing Multivariate Abundance Data.

Warton, D. I., Blanchet, F. G., O’Hara, R. B., Ovaskainen, O., Sara, T., Walker, S. C. & Hui, F. K. C. (2015). So Many Variables: Joint Modeling in Community Ecology. Trends in Ecology & Evolution, 30, 766 – 779.

Warton, D. I. & Hui, F. K. C. (2017). The central role of mean-variance relationships in the analysis of multivariate abundance data: a response to Roberts (2017). Methods in Ecology and Evolution, 8, 1408–1414.

Warton, D. I., Thibaut, L. & Wang, Y. A. (2017). The PIT-trap — A “model-free” bootstrap procedure for inference about regression models with discrete, multivariate responses. PLOS ONE, 12, 1–18.

Wickham, H., Cook, D., Hofmann, H. & Buja, A. (2011). tourr: An R Package for Exploring Multivariate Data with Projections. Journal of Statistical Software, 40, 118.

Wood, S. N. (2017). Generalized additive models: an introduction with R. CRC press.

Zuur, A. F., Ieno, E. N. & Smith, G. M. (2007). Principal component analysis and redundancy analysis. Analysing ecological data, pp. 193–224.

